# Epidemic potential of *Maravalia cryptostegiae*, a classical biological control agent of rubber vine, in Northeastern Brazil

**DOI:** 10.64898/2026.10.05.756685

**Authors:** Emerson M. Del Ponte, Rafael C. de Costa, Robert H. Barreto

## Abstract

Rubber vines (*Cryptostegia madagascariensis* and *C. grandiflora*) are highly invasive weeds in the semi-arid landscapes of many tropical regions. The rust fungus *Maravalia cryptostegiae* has provided a spetatular and sustainable control of invasive *C. grandiflora* populations in Australia, after its scientific and quarantine-authorized release in the context of a classical biological control program. The Caatinga biome of Northeastern Brazil is under a devastating invasion by *C. madagascariensis*. This species is also susceptible to *M. cryptostegia* but its epidemic potential towards *C. madagascariensis* in Brazil remains unknown. We developed a weather-driven epidemiological model integrating experimentally derived relationships for infection as a function of temperature and wetness duration and for temperature-dependent latent development. Three infection-response models were fitted to 20 experimental temperature × wetness combinations from the literature and compared using information criteria, grouped cross-validation and a parametric bootstrap. The selected model, in which thermal breadth increases with wetness duration, was best supported (R^2^ = 0.931; ΔAICc ≥ 3.3 relative to the alternatives), and prolonged wetness significantly broadened the thermal range favorable for infection (γ = 0.30 °C h^−1^; bootstrap P = 0.002). The infection and latent-development models were coupled with 25 years (2001–2025) of hourly NASA POWER data for 65 grid cells representing rubber-vine-infested locations in Northeastern Brazil, with wetness defined as hours with relative humidity ≥ 90%. Infection potential was concentrated in March–June (73% of the annual total, peaking in April), coinciding with the regional rainy season, and was highest in the interior of northern and central Ceará. Predicted potential varied more than a hundredfold among grid cells and was lowest along parts of the Ceará coast and the lower Jaguaribe valley, which include some of the largest infestations; the magnitude of these contrasts depended on the humidity threshold used to define wetness. Applied with identical rules to 13 historical release sites in Queensland, Australia, the model reproduced the documented wet-season timing of rust activity, and predicted potential was on average higher at sites with reported high rust impact, although the difference was not statistically significant. These results indicate strong seasonal and geographic structuring of weather-driven epidemic potential and provide testable hypotheses for selecting release periods and locations for *M. cryptostegiae* in Brazil.

## Introduction

Invasive alien plants are major drivers of ecological change and can impose substantial loss of biodiversity as well as harm ecosystem services, and agricultural production (Charles and Dukes 2007; Hejda et al. 2009). These impacts can be particularly difficult to manage in semi-arid landscapes, where invasive plants may occupy extensive and poorly accessible remote areas and conventional control measures are both economically unsustainable and ecologically harmful. In Northeastern Brazil, rubber vine (*Cryptostegia madagascariensis*: Apocynaceae), a woody climbing species native to Madagascar, has become a devastating invasive plant in the Caatinga biome (Seier et al. 2023). Surveys have documented *Cryptostegia* infestations across multiple Brazilian states and environments in that region, including riparian and seasonally flooded habitats where dense infestations can interfere with native vegetation and land use (Bonilla et al. 2023). Water loss through rubber vine evapotranspiration is also a major cause of concern in drought-prone areas of Northeastern Brazil.

Classical biological control offers a potentially sustainable approach for suppressing *Cryptostegia* over large and environmentally heterogeneous landscapes. A compelling precedent comes from northern Australia, where the autoecious rust fungus *Maravalia cryptostegiae* was introduced from Madagascar in 1995 for classical biological control of *C. grandiflora*. Following releases at 69 sites between 1995 and 1997, the rust fungus became established throughout much of the weed’s invaded range in Queensland and dispersed more than 500 km within three years (Tomley and Evans 2004). Recurrent rust epidemics caused extensive defoliation and substantially reduced plant biomass and fecundity; at some sites, the combined effects of rust and drought were associated with mortality of up to 75% of established plants, while seedling recruitment became nearly absent (Tomley and Evans 2004). These outcomes demonstrate the capacity of *M. cryptostegiae* to exert strong population-level effects on rubber vine, but they also highlight an important feature of the system: rust activity and impact vary markedly in space and time.

Such variability is expected because successful epidemics of foliar fungal pathogens depend not simply on broad climatic suitability, but also on the occurrence and temporal sequence of weather conditions that permit infection, particularly suitable combinations of temperature and leaf wetness duration (Duthie 1997; Magarey et al. 2005). For many foliar pathogens, infection is strongly determined by the joint effects of temperature and the duration of continuous leaf wetness, whereas temperature also regulates the rate at which latent infections develop toward sporulation (Duthie 1997). Consequently, two locations with similar seasonal mean temperature or rainfall may differ substantially in epidemic potential if they differ in the frequency, duration, and temperature of individual wetness episodes. Broad-scale climatic niche and species distribution models, including approaches such as CLIMEX and MaxEnt, are useful for estimating potential geographic distributions and environmental suitability (Phillips et al. 2006; Kriticos et al. 2024), but they typically do not resolve the short-term weather sequences governing infection and subsequent epidemic development.

Temperature–wetness response models provide a means of scaling experimentally determined infection biology to weather-driven estimates of infection potential. Controlled-environment studies commonly quantify infection over factorial combinations of temperature and wetness duration and fit nonlinear response surfaces to these data (Duthie 1997; Furuya et al. 2009; Arauz et al. 2010). These approaches have been used to estimate relative infection and construct weather-based infection-risk functions for diverse foliar pathogens. Importantly, temperature and wetness need not act as independent or separable drivers. The shape of the thermal response itself may change with wetness duration. For example, models developed for rust infection have shown improved fit when parameters controlling the temperature-response curve were allowed to vary with wetness duration (Furuya et al. 2009). Thus, explicitly representing temperature × wetness interactions may be necessary when prolonged wetness changes not only the magnitude of infection but also the range of temperatures over which infection can occur.

This interaction appears particularly relevant for *M. cryptostegiae*. Controlled-environment experiments by Evans and Fleureau (1993) quantified rust infection across combinations of temperature and dew duration and showed that infection was strongly constrained under short dew periods, whereas prolonged wetness supported substantial infection across a broader temperature range. The same study demonstrated a strong temperature dependence of latent development. Together, these experimentally quantified relationships provided an unusual opportunity to construct an epidemiologically explicit model in which hourly weather is translated sequentially into wetness episodes, relative infection potential, and latent development. Such a framework differs conceptually from a climatic suitability model because it represents the weather-dependent processes through which epidemics can develop.

Here, we developed a weather-driven epidemiological framework to quantify the seasonal and geographic epidemic potential of *M. cryptostegiae* at rubber-vine-infested locations in Northeastern Brazil. We first compared alternative nonlinear infection models, representing increasingly explicit forms of temperature–wetness interaction, ranging from a separable temperature × wetness response to models in which wetness duration modifies the thermal response. We then coupled the selected infection function with a temperature-dependent latent-development model and 25 years of hourly meteorological data to reconstruct infection opportunities and potential epidemic development across the invaded region. Finally, we evaluated the transferability of the framework using historical field observations from Queensland, Australia, where the rust has been established as a successful biological control agent for three decades.

Specifically, we addressed three questions: (i) does prolonged wetness alter the thermal breadth of *M. cryptostegiae* infection sufficiently to justify an explicit temperature × wetness interaction; (ii) when and where do weather conditions at known *Cryptostegia*-infested sites in Northeastern Brazil support infection and repeated epidemic development; and (iii) does the resulting weather-driven framework reproduce the contrasting seasonal and geographic patterns of rust activity documented in Queensland? By linking experimentally quantified infection biology with hourly weather, our aim was to move from broad climatic suitability toward a process-informed assessment of when and where *M. cryptostegiae* may have the greatest potential to develop epidemics following release.

## Material and Methods

Experimental observations of rust pustule density (*pustules cm*^−2^) under controlled combinations of temperature (*T*∈{17, 20, 22, 25, 30} °*C*) and continuous dew duration (*W*∈{6, 8, 12, 24} *h*) were obtained from Evans and Fleureau (1993). The complete experimental dataset comprises *N* = 20 treatment combinations (5 *temperatures* × 4 *wetness durations*). Testing *W* = 8 *h* wetness is physiologically critical because it captures the transition threshold between minimal infection initiation (*W* = 6 *h*, yielding ≤ 1% relative infection) and high-intensity infection saturation (*W*≥12 *h*). Maximum observed pustule density was 14. 35 *pustules cm*^−2^ at *T* = 22°*C* and *W* = 24 *h*. Relative infection response *I*_*obs*_ was normalized as:

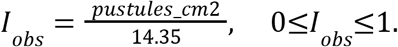

Experimental latent period observations (*LP*, days; *N* = 10) were reconstructed across temperatures *T*∈[18, 27] °*C*, and development rate was defined as *r* = 1/*LP* (*day* ^−1^). We evaluated three primary non-linear response surface models representing relative infection potential *I*(*T, W*) ∈ [0, 1], ordered by biological complexity:

### Model 1: Separable Generalized Beta × Weibull Model (Reference Model)

This model assumes complete separability between temperature and wetness duration, where temperature determines the unimodal thermal response curve and wetness duration scales infection magnitude (Hau and Kranz 1990):

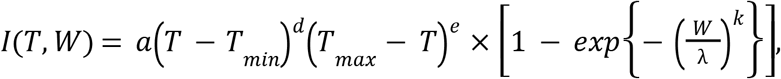

where lower (*T* _*min*_ = 10^°^ *C*) and upper (*T* _*max*_ = 35 °*C*) cardinal temperatures are fixed based on fungal biology, *a* is a scaling constant, *d* and *e* are shape parameters regulating thermal curve skewness, and λ and *k* are Weibull wetness scale and shape parameters. Model 1 contains 5 estimated parameters (*a, d, e*, λ, *k*).

### Model 2: Wetness-Expanded Analytis Beta × Weibull Model (General Interaction Comparator)

This other model serves as a general interaction comparator, allowing wetness duration to modify the temperature response curve without assuming specifically that modification occurs through Gaussian thermal breadth (Analytis 1977):

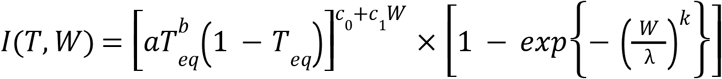

where 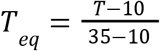 is equivalent temperature, *a* controls amplitude, *b* governs skewness, and *c*(*W*) = *c*_0_+ *c*_1_ *W* allows the shape exponent regulating curve width to expand linearly with wetness duration. Model 2 contains 6 estimated parameters (*a, b, c*_0_, *c*_1_, λ, *k*).

### Model 3: Gaussian Temperature Response with Centered Wetness-Dependent Breadth × Weibull Model (Selected Model)

Finally, this model represents a specific biological interaction in which prolonged wetness duration linearly expands the thermal breadth (σ) of the infection curve:

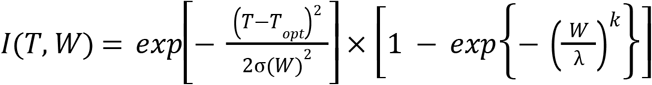

To improve parameter estimation, avoid ill-conditioned parameter correlations, and enforce a lower physiological bound on thermal breadth, thermal breadth was reparameterized using a centered wetness formulation:

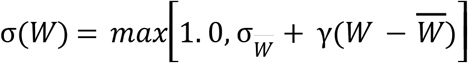

where *T*_*opt*_ is optimum temperature (°*C*), 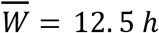 is the mean wetness duration across experimental observations (*W*∈{6, 8, 12, 24} *h*), 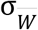 is thermal breadth at mean wetness duration (°*C*), γ is thermal breadth expansion per additional hour of wetness (^°^*C h*^−1^), and λ and *k* are Weibull wetness parameters. Model 3 contains 5 estimated parameters (*T*_*opt*_, σ_*W*−_, γ, λ, *k*).

### Supplementary Historical Models

For sensitivity and historical comparison, two additional models were evaluated (see Supplementary Material): Simple Analytis Beta × Weibull (*c*_1_ = 0; Model S1) and the Duthie Hyperbolic Secant × Weibull model [Duthie (1997); Model S2].

### Model Fitting & Goodness-of-Fit Comparison

All models were fitted to exactly the same observations (*N* = 20) and response scale (*I* _*obs*_) using robust non-linear least squares (nlsLM package in R, Levenberg-Marquardt algorithm) combined with multistart optimization grids across biologically plausible parameter ranges. Goodness-of-fit was evaluated using Root Mean Square Error (*RMSE*), coefficient of determination (*R*^2^, descriptive measure), log-likelihood (*logL*), Akaike Information Criterion (*AIC*), and corrected Akaike Information Criterion (*AICc*):

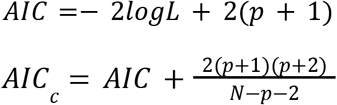

where *RSS* is residual sum of squares and *p* is the number of estimated regression parameters (one is added for the residual variance), and logL is the Gaussian log-likelihood at the least-squares solution. *AICc* served as the principal selection criterion due to the small sample size (*N* = 20). Model evidence was expressed as Δ*AICc* = *AICc* _*i*_ − *AICc* _*min*_.

### Model Validation via Grouped Cross-Validation

To assess interpolation and extrapolation performance across experimental conditions, we implemented two grouped cross-validation strategies:

Leave-One-Wetness-Level-Out Cross-Validation (LOWO): For each wetness duration *w*∈{6, 8, 12, 24} *h*, all 5 observations at that wetness level were omitted (training *N* = 15), the model was refitted, and predictions were evaluated on the omitted wetness level. Pooled prediction errors yielded LOWO RMSE.. Leave-One-Temperature-Level-Out Cross-Validation (LOTO): For each temperature level *t*∈{17, 20, 22, 25, 30} °*C*, all 4 observations at that temperature were omitted (training *N* = 16), the model was refitted, and predictions were evaluated on the omitted temperature. Pooled prediction errors yielded *LOTO RMSE*. In every training fold, models were refitted from 12 starting values (the default start, the full-data solution and 10 randomly perturbed starts), retaining the solution with the lowest residual sum of squares.

### Formal Evaluation of Wetness-Dependent Thermal Breadth (γ)

The biological parameter of interest is γ, measuring change in thermal breadth per hour of wetness. The null hypothesis of fixed thermal breadth is: *H*_0_ : γ = 0 versus the alternative of dynamic thermal expansion: *H*_*A*_ : γ > 0. Because γ≥0 lies on the boundary of the parameter space, standard likelihood-ratio tests, using a naive 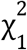 distribution, are invalid. Therefore, we performed a parametric bootstrap under *H*_0_ (*B* = 2, 000 replicates): 1. Fit the null fixed-breadth model (*H*_0_ : Model 3 with γ = 0, *p* = 4); 2. Generate *B* = 2, 000 simulated datasets under *H*_0_ by adding normal errors 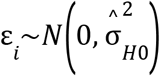 to *H*_0_ fitted values; 3. Refit both *H*_0_ (fixed) and *H*_1_ (dynamic) models to each simulated dataset, starting the dynamic-breadth fit from the fixed-breadth solution (γ = 0) and from additional starting values and retaining the lowest residual sum of squares, which guarantees ΔRSS ≥ 0; 4. Calculate the test improvement statistic Δ*RSS* = *RSS*_*H*0_ − *RSS*_*H*1_ ; 5. Calculate the bootstrap *P*-value as (x + 1)/(B + 1), where x is the number of simulated replicates in which Δ*RSS* _*b*_ ≥Δ*RSS*_*obs*_.

### Latent Development Submodel & Digitization Sensitivity

Primary linear rate regression *r*(*T*) = *a* + *bT* was compared against a non-linear quadratic rate model *r*(*T*) = *a* + *bT* + *cT*^2^ . Performance was evaluated on both the development rate scale (*R*^2^, *RMSE* _*rate*_) and back-transformed latent period scale (*RMSE*_*LP*_, days). To evaluate model sensitivity to graphical extraction precision, *B* = 500 stochastic iterations were executed by adding uniform noise (± 0. 05 *pustules cm*^−2^ for infection; ± 0. 5 *days* for latent period) around digitized points. Parameter stability percentiles (5th, 50th, 95th) and digitization sensitivity bands were derived.

### Hourly Meteorological Dataset and Epidemic Potential Calculation

Hourly meteorological data (2001–2025; 25 complete years) were retrieved using the nasapower R package (Sparks 2018) for 65 grid cells covering host distribution areas in Northeastern Brazil (14.2 million hourly records). An hour was classified as wet when relative humidity at 2 m was ≥ 90% (thresholds of 85% and 95% were evaluated as sensitivity analyses), and each continuous run of wet hours defined a wetness episode *j*. Precipitation was not used in the primary definition because hourly NASA POWER precipitation (PRECTOTCORR) was > 0 mm h^−1^ in 50–74% of all hours at the study grid cells, which would merge consecutive days into multi-day episodes; a variant in which hours with precipitation ≥ 1 mm h^−1^ were also classified as wet was evaluated as a sensitivity analysis. For each episode *j* with duration *W*_j_ ≥ 6 h and mean temperature *T*_wet,j_, relative infection potential *I*_j_ = *I*(*T*_wet,j_, *W*_j_) was calculated using Model 3. To avoid extrapolating the wetness-dependent expansion of thermal breadth beyond the experimental range, σ(*W*) was held at σ(24 h) for episodes longer than 24 h. For each cohort *j* (*I*_j_ > 0), hourly latent development accumulated from *t*_init,j_ = *t*_end,j_ + 1:

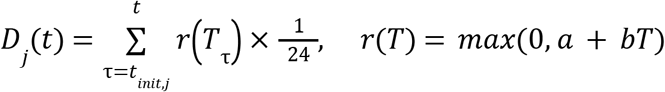

Hourly temperatures above 27 °C, the upper limit of the latent-period data, were set to 27 °C when computing *r*(*T*). The latent period of cohort *j* was defined as the time until *D*_j_(*t*) ≥ 1.0. Monthly and annual site-level epidemic potential was quantified as the infection pressure index, EPI_inf_ = Σ_j_ *I*_j_, i.e., the summed relative infection potential of all wetness episodes in the period. A sporulation-weighted index was not used because virtually all cohorts completed latent development under the observed temperatures, so it would duplicate EPI_inf_. Spatial variation across Northeastern Brazil was mapped and overlaid with the 73 municipal host infestation sites (*C. madagascariensis*) reported by Bonilla et al. (2023). Annual indices were averaged over all 25 years, with years (or months) without infection contributing zero.

### Australian Field Model Validation

To evaluate cross-continental generalizability under open field conditions, we extracted historical release and 10-year monitoring data from tropical and central Queensland, Australia (Tomley and Evans 2004). The Australian dataset comprises 13 strategic release and monitoring locations covering diverse climatic zones (coastal moist airflow, riparian corridors, Gulf savannas, and dry inland semi-arid rangelands; Table 4).

For each Australian site, hourly weather data (2001–2025; 2.85 million records) were retrieved from NASA POWER and processed with exactly the same wetness definition, infection and latent-development functions, and code as the Brazilian sites. The model was compared with two field benchmarks reported by Tomley and Evans (2004): (1) seasonal timing, i.e., whether predicted infection pressure (EPI_inf_) peaks during the Southern Hemisphere summer wet season; and (2) spatial contrast, i.e., whether predicted mean annual EPI_inf_ was higher at the five sites where severe rust damage, defoliation or weed mortality was reported (McLeod River, Laura Lea, Lakefield NP, Charters Towers and Inkerman) than at the remaining eight sites, for which spread or moderate rust was reported. Because the field observations are heterogeneous and do not coincide with the weather record (field monitoring 1995–2004; weather 2001–2025), the second comparison was treated as descriptive and summarized with Welch’s *t*-test and the Mann–Whitney test.

## Results

### Observed Biological Infection Pattern

In experimental infection trials (Evans and Fleureau 1993), rust pustule density responded strongly to both temperature and wetness duration (Figure 1A, B). Infection was minimal at *W* = 6 *h* across all temperatures (≤ 0. 65 *pustules cm* ^−2^; *I* _*obs*_≤0. 045), indicating that 6 *h* represents the lower physiological threshold for infection initiation. At *W* = 8 *h*, infection increased substantially, peaking at 22 ^°^ *C* (6. 35 *pustules cm* ^−2^; *I* _*obs*_ = 0. 442). At extended wetness durations (*W* = 12 *h* and *W* = 24 *h*), infection saturated near optimal temperatures (12. 95 − 14. 35 *pustules cm* _−2_; *Iobs* = 0. 903 − 1. 000) and, crucially, high infection levels expanded into cooler (17 °*C*) and warmer (25°*C*) temperatures. This biological pattern indicates that prolonged wetness duration not only increases infection intensity but also broadens the thermal window favorable for rust infection.

**Figure 1.**
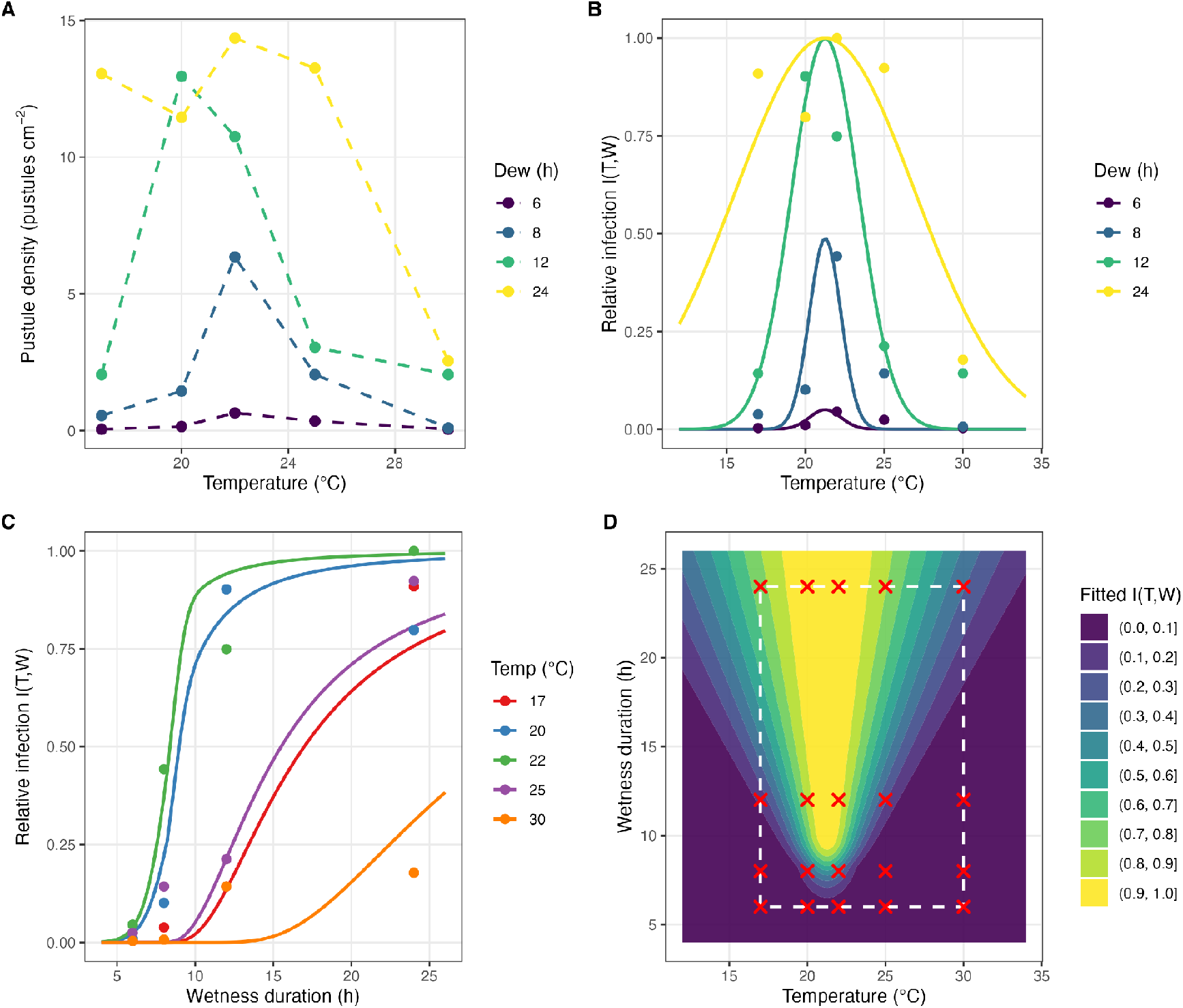
Selected Infection Submodel (Model 3) 2 × 2 Panel Grid. (A) Raw observed rust pustule density (*pustules cm* ^−2^) across temperatures for each wetness duration (6, 8, 12, 24 *h*); (B) Model 3 fitted infection potential (*I*(*T, W*)) across temperature slices (6, 8, 12, 24 *h*); (C) Model 3 fitted infection potential across wetness duration slices (17, 20, 22, 25, 30 °*C*); (D) Continuous response surface contour plot of fitted *I*(*T, W*) with white dashed experimental support rectangle and red crosses marking treatment combinations.xs

### Infection Model Comparison and Evidence Evaluation

All three primary infection submodels converged (Table 1). Model 3 (Gaussian temperature response with wetness-dependent breadth × Weibull) had the lowest AICc (R^2^ = 0.931, RMSE = 0.0973). Model 2, the general interaction comparator, fitted the data almost as closely (R^2^ = 0.936, RMSE = 0.0935) but with one additional parameter and was less supported (ΔAICc = 3.29). The separable reference Model 1 was clearly inferior (ΔAICc = 17.08), as were the historical Duthie model and the fixed-breadth Gaussian model (ΔAICc = 13.60 and 13.11, respectively; Table S1).

**Table 1.** Goodness-of-fit metrics comparing primary candidate infection submodels: Model 1 (Separable Generalized Beta × Weibull), Model 2 (Wetness-Expanded Analytis Beta × Weibull), and Model 3 (Gaussian Wetness-Dependent Thermal Breadth × Weibull).

| Model | Parameter<sup>a</sup> | RMSE<sup>b</sup> | $R^2$<sup>c</sup> | LogLik<sup>d</sup> | AIC<sup>e</sup> | AICc<sup>f</sup> | $\Delta\text{AICc}$<sup>g</sup> |
| --- | --- | --- | --- | --- | --- | --- | --- |
| Model 1 | 5 | 0.1491 | 0.8379 | 9.6795 | -7.3591 | -0.8975 | 17.0795 |
| Model 2 | 6 | 0.0935 | 0.9362 | 19.0121 | -24.0243 | -14.6910 | 3.2861 |
| Model 3 | 5 | 0.0973 | 0.9310 | 18.2193 | -24.4386 | -17.9771 | 0.0000 |

<sup>a</sup> number of estimated model parameters; <sup>b</sup> root mean square error; <sup>c</sup> coefficient of determination; <sup>d</sup> log-likelihood
<sup>e</sup> Akaike information criterion; <sup>f</sup> Akaike information criterion corrected for small sample size; <sup>g</sup> $\Delta\text{AICc}$ , difference in AICc relative to the model with the lowest AICc. $\text{AIC} = -2\log L + 2(p + 1)$ , with one parameter added for the residual variance.

Grouped cross-validation gave qualified support for Model 3. Model 3 had the lowest pooled prediction error both when entire wetness levels were withheld (LOWO RMSE = 0.237, compared with 0.339 for Model 2, 0.316 for the fixed-breadth model and 0.849 for Model 1) and when entire temperature levels were withheld (LOTO RMSE = 0.170, compared with 0.225, 0.189 and 0.200, respectively). Its advantage in LOWO arose mainly from the 24-h fold: when the 24-h treatment was withheld, Model 3 predicted it with RMSE = 0.137, whereas all other models failed to reproduce the broad thermal response observed under prolonged wetness (RMSE = 0.45–1.64). In the remaining folds, simpler models performed similarly or better, and predictions for withheld wetness levels were sensitive to starting values because the Weibull wetness term was poorly constrained when only three wetness levels were available for training. Cross-validation is therefore interpreted as supporting rather than decisive evidence for Model 3.

Parameter estimates for Model 3 (Table 2) identified an optimum infection temperature of T_opt_ = 21.26 ± 0.21 °C. Thermal breadth at mean experimental wetness duration (*W*− = 12. 5 *h*) was σ(W) = 2.28 ± 0.25 °C. The thermal breadth expansion rate was γ = 0.298 ± 0.063 °C h^−1^ (t = 4.73, P < 0.001), indicating that each additional hour of wetness expands the thermal breadth by 0. 30 °*C*. Calculated thermal breadth was strictly positive and bounded across all experimental wetness durations: σ(6 *h*) = 1. 00 °*C*, σ_8 h_ = 1.00 °C, σ_12 h_ = 2.13 °C, and σ_24 h_ = 5.71 °C. At 6 and 8 h, σ(W) was at its lower bound of 1.0 °C, so γ is informed mainly by the contrast between the 12- and 24-h treatments.

**Table 2.** Parameter estimates and associated uncertainty for the selected infection model (Model 3: centered Gaussian dynamic breadth × Weibull).

| Parameter<sup>a</sup> | Estimate | SE<sup>b</sup> | 95% CI<sup>c</sup> |
| --- | --- | --- | --- |
| $T_{opt}$ | 21.2597 | 0.2068 | 20.82–21.70 |
| $\sigma_{\bar{W}}$ | 2.2798 | 0.2538 | 1.74–2.82 |
| $\gamma$ | 0.2984 | 0.0630 | 0.164–0.433 |
| $\lambda$ | 8.3666 | 0.5559 | 7.18–9.55 |
| $\kappa$ | 8.9288 | 9.1384 | — |

<sup>a</sup> $T_{opt}$ – optimum temperature for infection (°C), $\sigma_{\bar{W}}$ – thermal-response breadth at the centered wetness duration (°C), $\gamma$ – change in thermal-response breadth per hour of wetness (°C h<sup>-1</sup>), $\lambda$ – Weibull wetness-scale parameter (h), and $\kappa$ – dimensionless Weibull shape parameter; <sup>b</sup> SE, standard error; <sup>c</sup> CI, confidence interval. The Weibull shape parameter $\kappa$ was poorly identified (large SE), reflecting the step-like increase in infection between 8 and 12 h; no confidence interval is given.

### Parametric Bootstrap Inference for γ

The parametric bootstrap under *H*_0_: γ = 0 (*B* = 2, 000 replicates) provided strong statistical evidence against a fixed thermal breadth (Figure S1). The observed improvement statistic was ΔRSS_obs_ = RSS_H0_ − RSS_H1_ = 0.4495 − 0.1894 = 0.2601. Across 2, 000 bootstrap replicates generated under the fixed-breadth null model, three replicates achieved a Δ*RSS* exceeding the observed value, yielding a parametric bootstrap *P*-value of P = 0.002 [(3 + 1)/(2,000 + 1)]. The 95th percentile of the null distribution of ΔRSS was 0.068.

### Latent Development Submodel

Temperature exhibited a marked linear relationship with fungal development rate *r*(*T*) =− 0. 07815 + 0. 007181*T* (Figure 2B). Model comparison favored the linear over the quadratic formulation by Δ*AICc* =+ 4. 27, demonstrating that higher-order thermal deceleration was not detectable within the experimental range (18–27 ^o^C). On the back-transformed latent period scale, the model exhibited high accuracy (*RMSE*_*LP*_ = 1. 82 *days* Figure 2C–D). The corresponding base temperature threshold was T_0_ = 10.88 ^o^C, establishing a thermal requirement of approximately 139.3 degree-days above T_0_ to complete the latent period.

**Figure 2.**
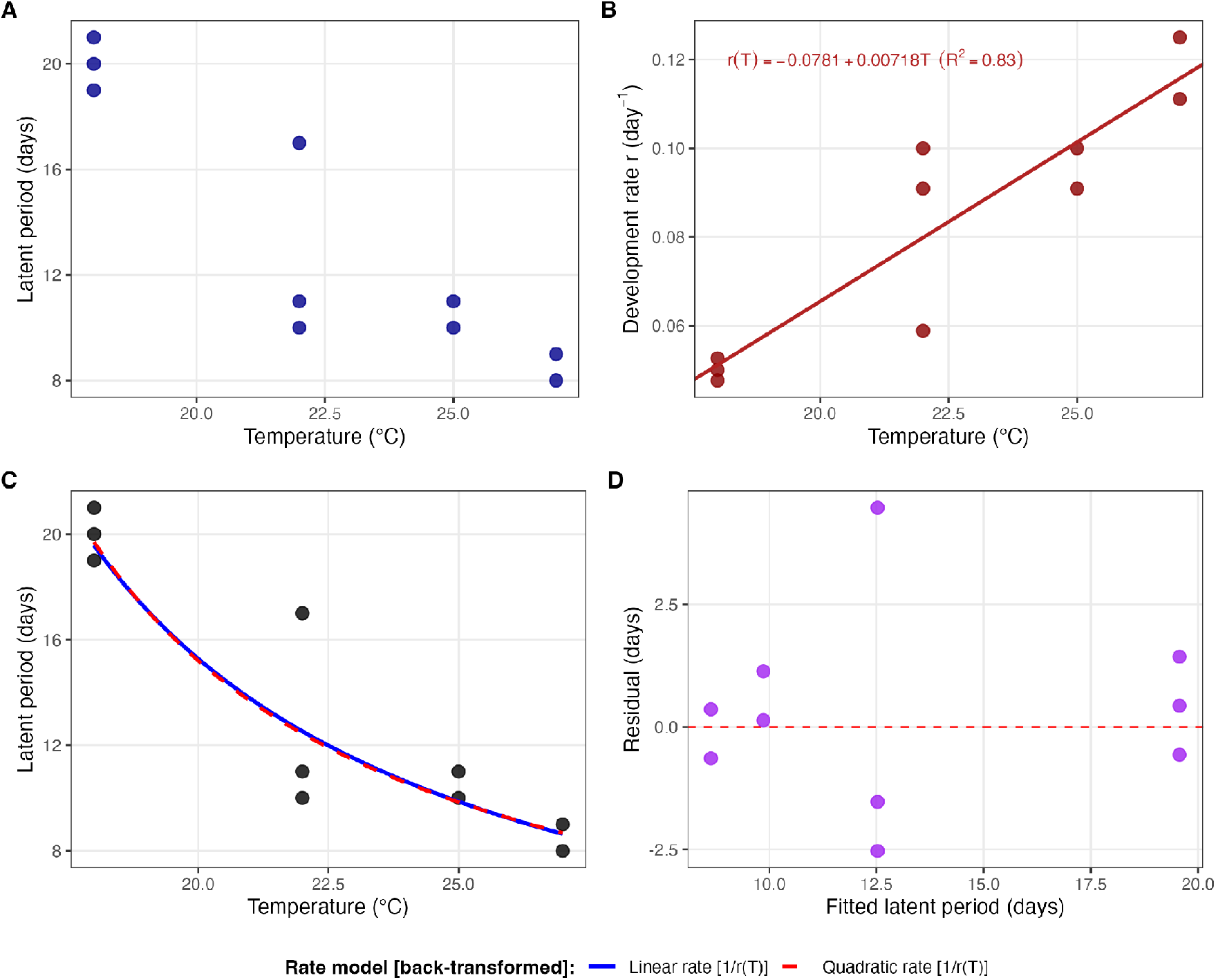
Effect of temperature on the latent period and development rate of *Maravalia cryptostegiae*. (A) Observed latent period (days) across constant incubation temperatures (18–27 ^o^C, N = 10). (B) Linear regression of fungal development rate (*r* = 1 / LP, in day ^-1^) as a function of temperature. Solid line denotes the ordinary least-squares fit; points represent observed rates. (C) Observed latent period (circles) and back-transformed predictions from the linear rate model (*LP* = 1 / *r*(*T*); solid blue curve) and the alternative quadratic rate model (dashed red curve). (D) Residual plot of the back-transformed primary linear model (observed minus fitted latent period, in days) plotted against fitted latent periods; the dashed red line represents zero residual error (*y* = 0).

### Digitization Sensitivity Analysis

Stochastic perturbation (*B* = 500 iterations) demonstrated high parameter stability under graphical extraction noise (Figure S4). *T*_*opt*_ varied by < 0. 02^°^ *C* (21.25–21.27 °C; 5th–95th percentiles), 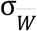 varied between 2.27 and 2.29 °C, and γ varied between 0.2965 and 0.3004 °C h^−1^ (Table S2).

### Wetness Episode Climatology in Northeastern Brazil

Under the primary wetness definition (RH ≥ 90%), 305,537 wetness episodes were identified across the 65 grid cells over the 25-year period (about 188 per cell and year; Table 3). Episodes lasted on average 9.4 h at a mean temperature of 23.5 °C, close to the infection optimum; 238,911 episodes (78.2%) lasted ≥ 6 h, 204,746 (67.0%) ≥ 8 h and 114,483 (37.5%) ≥ 12 h.

**Table 3.** Sensitivity of wetness and infection metrics to relative humidity thresholds.

| RH (%) | Episodes | $\geq 6$ h<sup>a</sup> | $\geq 8$ h | $\geq 12$ h | WD (h)<sup>b</sup> | $T_w$ (°C)<sup>c</sup> | Annual IP<sup>d</sup> |
| --- | --- | --- | --- | --- | --- | --- | --- |
| 85 | 409,038 | 319,077 | 281,498 | 183,202 | 10.21 | 23.85 | 86.75 |
| 90 | 305,537 | 238,911 | 204,746 | 114,483 | 9.41 | 23.48 | 62.83 |
| 95 | 198,793 | 129,036 | 90,418 | 22,912 | 7.02 | 22.96 | 25.35 |

<sup>a</sup> Episodes $\geq 6$ h, $\geq 8$ h, and $\geq 12$ h represent qualifying events for fungal infection; <sup>b</sup> mean duration of extracted continuous wetness episodes (hours); <sup>c</sup> mean air temperature during wetness (°C); <sup>d</sup> mean annual cumulative infection potential per site-year ( $EPI_{inf}$ ) across the 65 sites (2001–2025). Wet hours defined by relative humidity alone; $\sigma(W)$ held at its 24-h value for longer episodes.

Epidemic potential was sensitive to the humidity threshold used to define wetness (Table 3). Mean annual EPI_inf_ per grid cell was 86.7, 62.8 and 25.3 for thresholds of 85%, 90% and 95%, respectively, mainly because stricter thresholds shortened episodes (mean duration 10.2, 9.4 and 7.0 h) and sharply reduced the number of episodes lasting ≥ 12 h (183,202, 114,483 and 22,912). Additionally, classifying hours with precipitation ≥ 1 mm h^−1^ as wet reduced mean annual EPI_inf_ to 51.1 and shifted the monthly peak from April to May. Absolute values of the index should therefore be interpreted relative to the wetness definition.

### Latent Cohort Tracking and Seasonal Epidemic Window

Latent development was completed by 238,900 of 238,911 infection cohorts (> 99.9%), with a mean latent period of 9.8 days (range 8.7–12.5 days). Infection potential showed a distinct seasonal window (Figure 3A): EPI_inf_ increased from February, peaked in April (13.0 per month) and remained high through June, so that March–June accounted for 73% of the annual total, whereas September–December contributed < 1%. This window coincides with the main rainy season of northern Northeastern Brazil.

**Figure 3.**
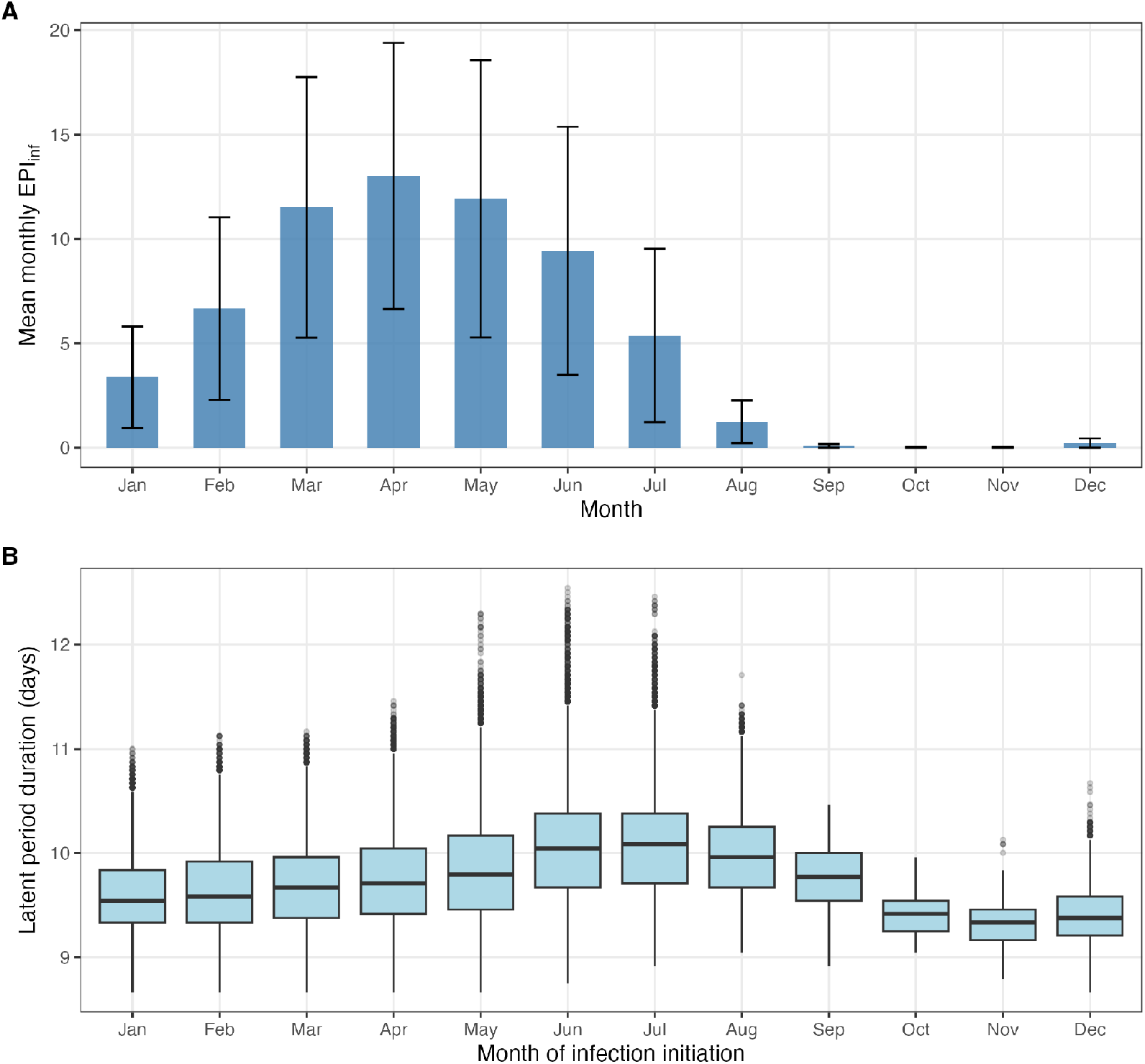
Monthly climatology across Northeastern Brazil (2001–2025). (A) Monthly infection pressure (*EPI*_*inf*_), mean ± SD across the 65 grid cells; (B) distribution of the latent period of infection cohorts by month of infection initiation.

### Spatial Risk Mapping & Host Infestation Overlay

Across the 73 infestation localities recorded by Bonilla et al. (2023), which fall in 28 distinct grid cells, mean annual EPI_inf_ averaged 62.9 but ranged from < 0.1 to 117.8 (Table S3). Potential was highest in the interior of northern and central Ceará (e.g., Varjota, Cariré, Sobral, Canindé and Itatira; EPI_inf_ = 105–118) and lowest in the lower Jaguaribe valley and on the coast west of Fortaleza (e.g., Jaguaruana, Aracati, Itaiçaba, Caucaia and São Gonçalo do Amarante; EPI_inf_ < 1), where RH ≥ 90% was rarely reached in the NASA POWER data (Figure 4A). Seven of the 73 localities had EPI_inf_ < 20. In western Paraíba (São João do Rio do Peixe, Sousa), EPI_inf_ was about 50.

**Figure 4.**
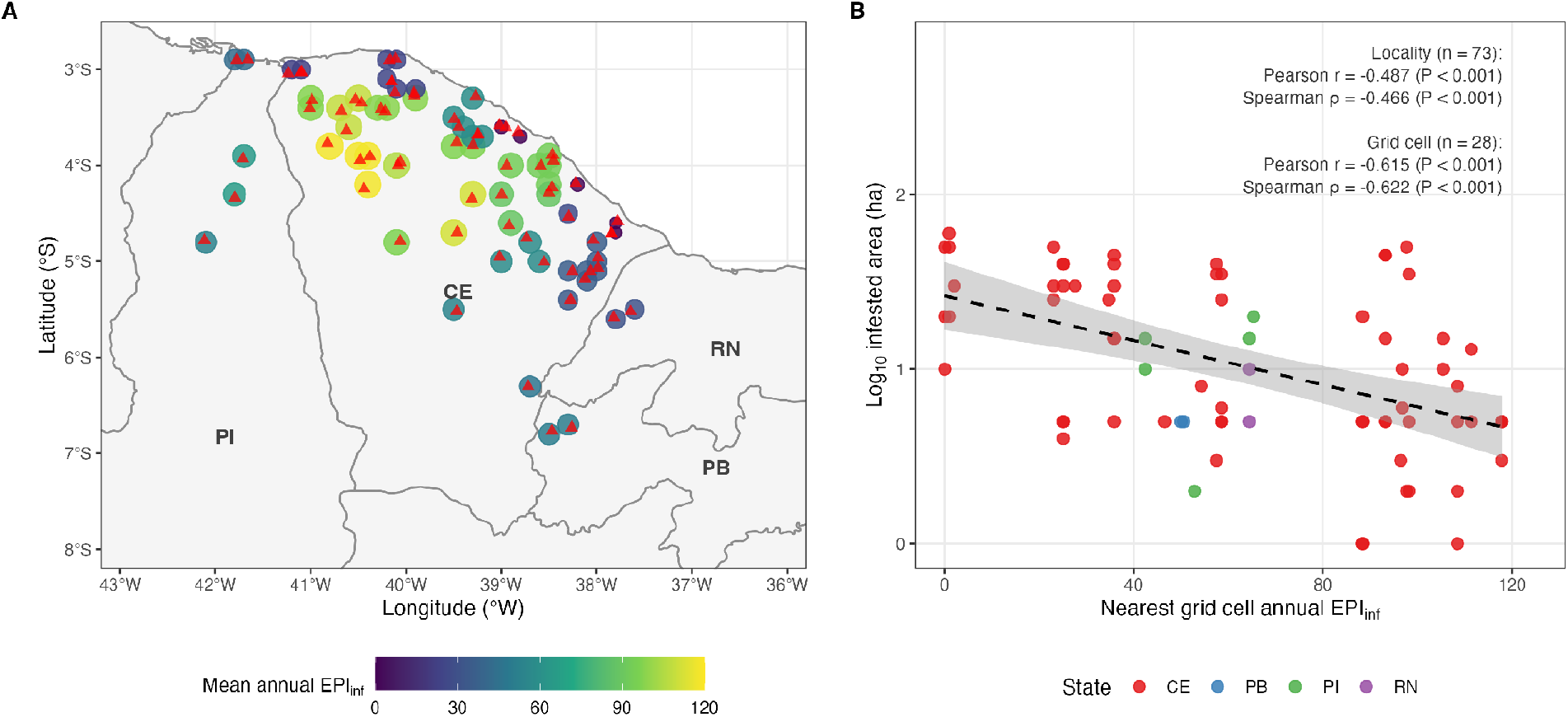
Spatial distribution of predicted infection pressure and host infestations in Northeastern Brazil. (A) Mean annual infection pressure (*EPI* _*inf*_) for each NASA POWER grid cell (circles; 2001–2025) and the 73 infestation localities reported by Bonilla et al. (2023) (triangles); (B) recorded host infestation area (*log*_10_ *ha*) vs. mean annual EPI_inf_ of the nearest grid cell, with linear trend and 95% confidence band; correlations are shown for localities (n = 73) and grid cells (n = 28).

Infested area was negatively related to predicted infection pressure across the 73 localities (log_10_ area vs. EPI_inf_: r = −0.49, P < 0.001; Spearman ρ = −0.47, P < 0.001; Figure 4B). Because localities sharing a grid cell have identical EPI_inf_, the association was also evaluated with one value per grid cell (mean log_10_ area; n = 28), giving ρ = −0.62 (P < 0.001). This relationship reflects the history of weed introduction and spread rather than any effect of the rust, which has not been introduced into Brazil. Several of the largest infestations (≥ 45 ha) lie in coastal and lower Jaguaribe municipalities with very low predicted potential (Caucaia, São Gonçalo do Amarante and Jaguaruana; EPI_inf_ < 1), whereas others (Granja, Aquiraz and Pacatuba; EPI_inf_ = 93–98) coincide with high potential. Interior municipalities with the highest potential currently harbor mostly small patches (1–15 ha).

### Australian Field Model Validation

Model validation across 13 historical field monitoring sites in Queensland, Australia (Tomley and Evans, 2004) was evaluated for seasonal timing and spatial contrast (Table 4, Figure 5).

**Table 4.** Cross-continental validation summary comparing predicted mean annual infection pressure (EPI_inf) and latent period against 10-year field release and impact monitoring observations across 13 sites in Queensland, Australia (Tomley & Evans 2004).

| Site | Lat. | Lon. | $EPI_{inf}$ | $LP_{days}$ | Observed field impact |
| --- | --- | --- | --- | --- | --- |
| McLeod River | -16.48 | 145.23 | 161.00 | 13.93 | Heavy rust (47 pustules cm<sup>2</sup>); total defoliation |
| Laura Lea | -15.56 | 144.44 | 80.09 | 10.81 | Enhanced rust intensity; 20% mortality by 1999 |
| Lakefield NP | -14.94 | 144.20 | 69.30 | 10.34 | High defoliation & 20-75% weed mortality |
| Chillagoe NP | -17.15 | 144.52 | 57.53 | 12.06 | Spread to 80 km by Aug 1996 |
| Charters Towers | -20.08 | 146.26 | 42.50 | 13.58 | Key monitoring hub; heavy rust in wet years |
| Wrotham Park | -16.63 | 144.00 | 41.44 | 10.68 | Rapid spread to 50 km in 8 months |
| Rockhampton | -23.38 | 150.51 | 35.83 | 14.64 | Rapid spread over 100 km in 14 months |
| Georgetown | -18.29 | 143.55 | 23.03 | 12.23 | Moderate rust; shorter wet season duration |
| Rutland Plains | -15.55 | 141.73 | 15.42 | 9.39 | Satellite spread >20 km detected |
| Strathmore | -17.85 | 142.37 | 11.68 | 9.65 | Spread to 45 km by Aug 1996 |
| Hughenden | -20.84 | 144.20 | 8.05 | 14.24 | Inadequate control in most years due to dry air |
| Inkerman | -19.74 | 147.46 | 5.74 | 10.95 | Severe rust damage; up to 75% mortality |
| Delta Downs | -17.10 | 140.92 | 0.72 | 10.31 | Spread to 70 km in 18 months |

**Figure 5.**
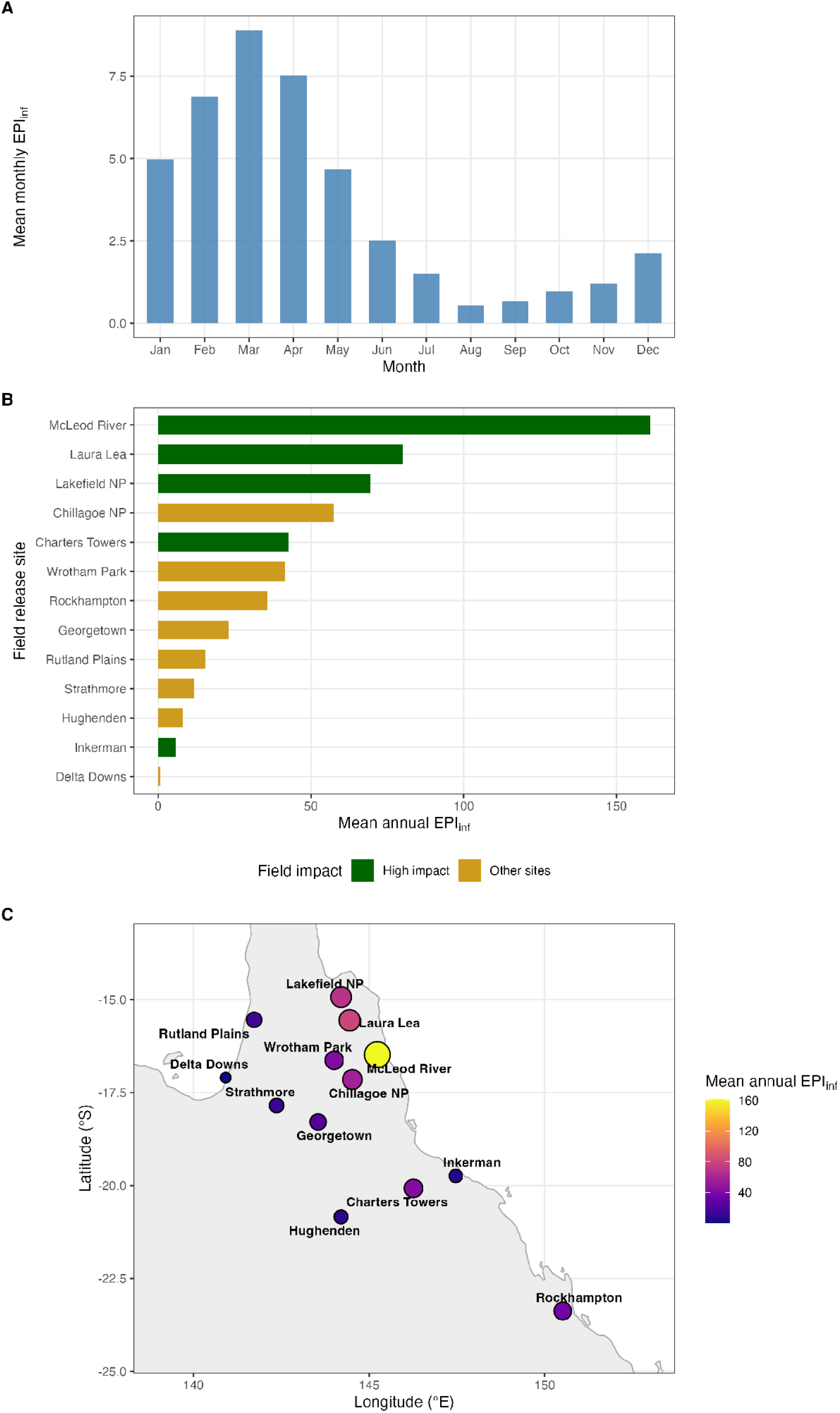
Model evaluation at Queensland release sites. (A) Monthly mean infection pressure (*EPI*_*inf*_) across 13 sites (2001–2025); (B) sites ranked by predicted mean annual EPI_inf_ and coloured by reported field impact (dark green = high impact, gold = other sites; Tomley and Evans 2004); (C) Spatial map of Queensland, Australia showing the 13 field release and monitoring locations colored and sized by predicted mean annual infection potential (*EPI* _*inf*_).

#### Seasonal Epidemic Climatology

In tropical Queensland (Southern Hemisphere), the model correctly predicted that weather-driven infection pressure (*EPI*_*inf*_) peaks during the summer wet season (January to April, peaking in March) (Figure 5A). This agrees with the wet-season timing of rust activity and releases reported by Tomley and Evans (2004).

#### Spatial Contrasts among Release Sites

Predicted potential varied widely among Queensland sites (EPI_inf_ = 0.7–161.0; Table 4). The three sites with the most severe documented impact had the highest values (McLeod River, 161.0; Laura Lea, 80.1; Lakefield NP, 69.3), and the dry inland sites where control was reported to be inadequate had low values (Georgetown, 23.0; Hughenden, 8.0). However, the correspondence was not consistent: Inkerman, where up to 75% weed mortality was recorded, had one of the lowest values (5.7), and rapid spread was documented at sites with low predicted potential (e.g., Delta Downs, 0.7; Strathmore, 11.7).

Mean annual EPI_inf_ at the five high-impact sites (71.7) was higher than at the remaining eight sites (24.2), but the difference was not statistically significant (Welch t = 1.78, df = 4.6, P = 0.14; Mann–Whitney W = 32, P = 0.093). With hours of precipitation ≥ 1 mm h^−1^ included in the wetness definition, the corresponding P-values were 0.087 and 0.045.

## Discussion

Our analysis indicates that weather across much of the area invaded by *C. madagascariensis* in northeastern Brazil supports recurrent infection and completion of the latent period of *M. cryptostegiae*, but that this potential is strongly seasonal and spatially uneven. Rather than describing climatic suitability in a general sense, our framework links hourly weather to two processes directly associated with epidemic development—infection and completion of the latent period. Opportunities for infection were concentrated in the rainy season (March–June) and were greatest in the interior of northern and central Ceará, whereas parts of the coast and the lower Jaguaribe valley, which include some of the largest infestations, showed little potential under the primary wetness definition.

A key feature emerging from the infection model is that temperature and wetness duration should not be considered independently. The experimental data of Evans and Fleureau (1993) indicate a relatively narrow thermal window for infection when wetness periods are short, whereas prolonged wetness permits substantial infection over a much broader temperature range. This interaction has important epidemiological consequences. Under marginal wetness duration, infection is largely restricted to periods when temperature is close to the optimum. Longer periods of surface moisture, in contrast, provide sufficient time for pre-penetration processes to proceed even when temperature is less favorable, thereby relaxing the thermal constraint on infection. Consequently, the occurrence and duration of wet periods may be at least as important as mean temperature in determining when epidemics can develop.

This behavior is consistent with the broader principle that temperature and wetness interact in determining foliar pathogen infection, although the form of that interaction varies considerably among pathosystems (Duthie 1997; Magarey et al. 2005; Monroe et al. 1997; Wu et al. 1999; Furuya et al. 2009; Arauz et al. 2010). Wetness duration may affect the amplitude, optimum, asymmetry, or thermal breadth of infection responses, and separable temperature and wetness functions are therefore not universally appropriate. For *M. cryptostegiae*, allowing thermal breadth to change with wetness duration provided a substantially better representation of the experimental response than models assuming a fixed thermal response. The parametric bootstrap indicated that this improvement is unlikely under a fixed-breadth model, and the wetness-dependent model was the only one that predicted the broad thermal response at 24 h when that treatment was withheld. Grouped cross-validation with only four wetness and five temperature levels was, however, unstable, so the interaction is best regarded as well supported by the available data rather than definitively established.

Nevertheless, the temperature–wetness function should be regarded as an empirical representation of the conditions tested by Evans and Fleureau (1993), rather than as a general physiological response function. In particular, the increase in thermal breadth with wetness duration is supported only within the experimental range. Extrapolating this relationship to substantially longer wet periods would eventually imply biologically unrealistic thermal responses. This limitation is especially important when transferring controlled-environment relationships to hourly weather data, where naturally occurring wet periods may extend beyond those represented experimentally. Restricting interpretation to the experimental domain, or explicitly evaluating alternative extrapolation rules, is therefore preferable to assuming indefinite expansion of the thermal window. In this study, σ(W) was therefore held at its 24-h value for longer episodes; with wetness defined by relative humidity, fewer than 0.1% of episodes exceeded 24 h, so this choice had a negligible effect on the results.

The comparison with the Australian experience provides an ecological benchmark for interpreting the Brazilian predictions. *Maravalia cryptostegiae* became an exceptionally effective biological control agent of rubber vine in Australia, where recurrent rust epidemics caused severe defoliation and substantial reductions in plant performance (Tomley and Evans 2004). When the same model, weather source and wetness definition were applied to both regions, predicted potential at most Brazilian host localities (median 58.6; three-quarters above the Queensland median of 35.8) was comparable to or higher than that at Queensland sites where the rust established and spread, and the Queensland wet-season peak was reproduced. The model did not, however, consistently separate high-impact from other Queensland sites: some sites with severe documented impact had low predicted potential, and the difference between site groups was not significant. These comparisons support the plausibility of weather-supported epidemics in much of the Brazilian invaded range but cannot establish that the rust will establish or achieve equivalent control.

An important aspect of this result is that the high epidemic potential predicted for much of interior northeastern Brazil is not inconsistent with the semiarid character of much of the region. Infection by foliar pathogens responds to the conditions experienced at the leaf surface over relatively short periods rather than to broad climatic classifications. Nocturnal cooling can produce prolonged periods of high relative humidity even where daytime atmospheric demand is high, while rainfall provides additional opportunities for extended wetness. Thus, environments conventionally characterized as dry may still contain recurrent nighttime windows suitable for rust infection. This illustrates why process-oriented analysis of hourly weather can reveal epidemiologically important conditions that would be obscured by seasonal rainfall or mean relative humidity alone. Conversely, the very low values predicted for some coastal and lowland cells should be interpreted cautiously, because gridded humidity may not represent near-surface conditions at the riparian and floodplain sites where rubber vine grows; comparison with station or leaf-wetness data is needed before these areas are considered unfavorable.

The strong seasonality of these infection opportunities also has practical implications for biological control. Releases should, in principle, be timed to coincide with the beginning of periods that combine frequent prolonged wetness with favorable temperatures, thereby increasing the probability that initial inoculum produces infection and is followed by additional weather-supported generations. In northeastern Brazil this corresponds to the onset of the rainy season (February–March), which would allow several infection cycles before conditions become unfavorable from August onwards. Similarly, geographic prioritization could focus initial releases on locations where large host patches coincide with high predicted potential (e.g., Granja, Aquiraz and Pacatuba). These predictions should be regarded as hypotheses for field evaluation rather than prescriptive release recommendations. Field releases across contrasting levels of predicted epidemic potential, including low-potential coastal localities, would provide a particularly powerful prospective test of the model.

Several sources of uncertainty remain. Leaf wetness is inferred from meteorological variables rather than measured directly at the host canopy, and relationships developed under continuous controlled dew periods necessarily simplify the fluctuating temperature, humidity, radiation, and wetness conditions experienced in the field. The infection and latent-development functions were also reconstructed from published experimental figures rather than fitted to the original observations. More fundamentally, the present framework represents the weather component of epidemic potential. It does not explicitly account for inoculum availability, host density and phenology, canopy microclimate, dispersal, loss of viable spores, or feedback between rust-induced defoliation and subsequent host availability. These processes may determine whether favorable meteorological conditions are translated into sustained epidemics and effective weed suppression. In particular, the absolute level of the index and its spatial contrasts depended strongly on the relative humidity threshold used to define wetness (mean annual EPI_inf_ of 25–87 for thresholds of 95–85%), and hourly NASA POWER precipitation proved unsuitable as a wetness indicator. Validating the wetness proxy against station humidity or leaf-wetness measurements in the Caatinga is therefore a priority.

Accordingly, the model should not be interpreted as predicting rust severity or guaranteeing establishment of *M. cryptostegiae*. Its principal contribution is to translate experimentally determined infection and development responses into a spatially and temporally explicit measure of weather-driven epidemic potential. Within that scope, the convergence between the experimental biology of the pathogen, the reconstructed Australian experience, and the climatic conditions encountered across invaded areas of northeastern Brazil provides an epidemiological basis for field evaluation of *M. cryptostegiae* as a biological control agent of *C. madagascariensis*.

---

**Table S1.**
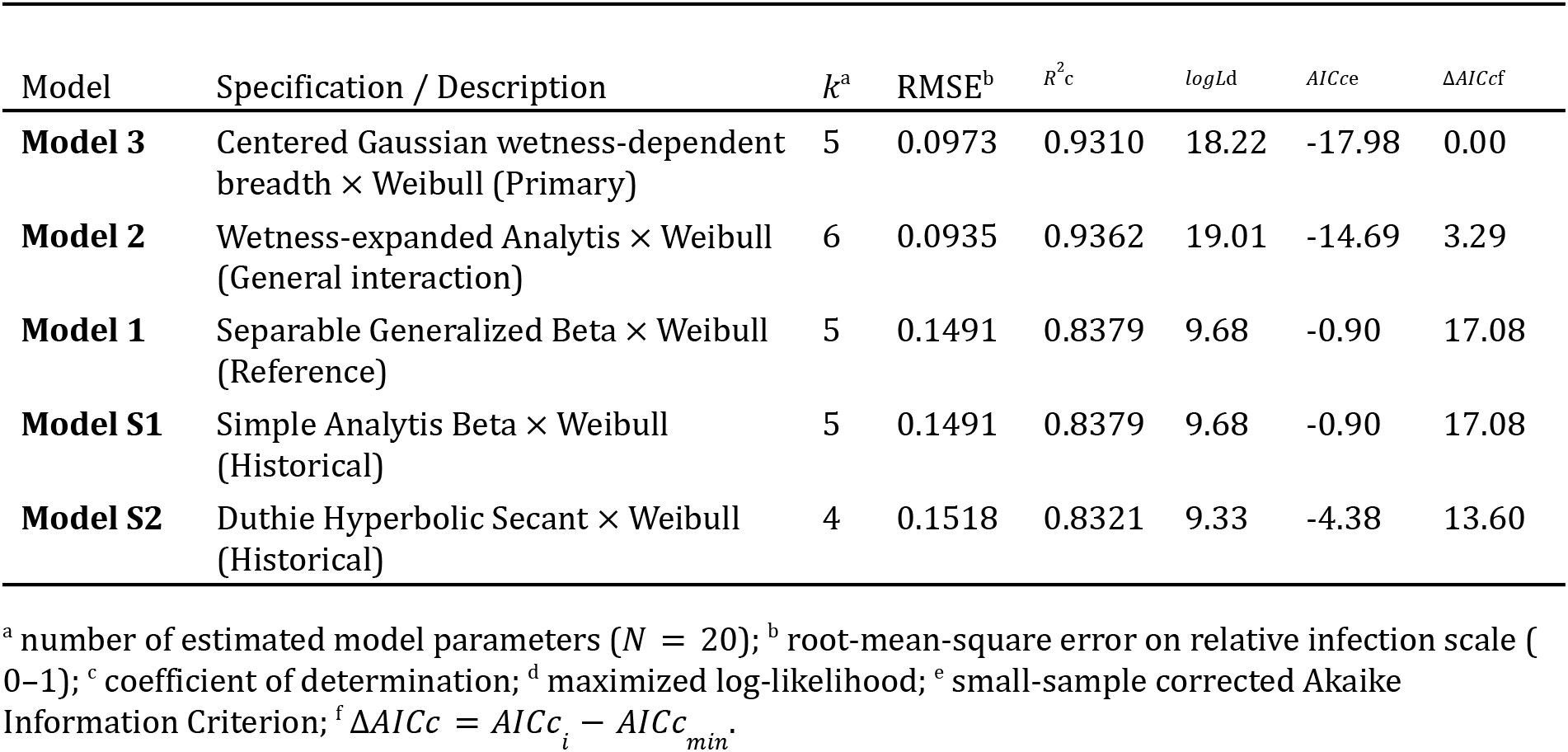
Information-theoretic comparison and goodness-of-fit statistics across all candidate infection submodels for Maravalia cryptostegiae. Models 1–3 represent the primary candidate set; Models S1 and S2 represent historical epidemiological formulations; Model S3 represents the nested fixed-breadth null hypothesis evaluated in the parametric bootstrap.

**Table S2.** Parameter stability and sensitivity analysis of infection and latent development submodels under stochastic graphical digitization error (*B* = 500 iterations). Uniform random perturbations (± 0. 05 *pustules cm*^−2^ for infection intensity; ± 0. 5 *days* for latent period) were introduced around digitized data points.

| Submodel | Parameter | Baseline Estimate | 5th Percentile | Median (50th) | 95th Percentile |
| --- | --- | --- | --- | --- | --- |
| Infection Submodel | $T_{opt}$ (optimum temperature, $^{\circ}C$) | 21.26 | 21.25 | 21.26 | 21.27 |
| | $\sigma_{\overline{W}}$ (thermal breadth at $\overline{W} = 12.5$ *h*, $^{\circ}C$) | 2.28 | 2.27 | 2.28 | 2.29 |
| | $\gamma$ (thermal breadth expansion rate, $^{\circ}C$ *h*$^{-1}$) | 0.2984 | 0.2965 | 0.2984 | 0.3004 |
| | $\lambda$ (Weibull scale parameter, *h*) | 8.37 | 8.35 | 8.37 | 8.38 |
| | $k$ (Weibull shape parameter, dimensionless) | 8.93 | 8.69 | 8.95 | 9.20 |
| Latent Development | $a$ (rate intercept, *day*$^{-1}$) | -0.0781 | -0.0866 | -0.0788 | -0.0710 |
| | $b$ (rate slope, *day*$^{-1}$ $^{\circ}C$ $^{-1}$) | 0.00718 | 0.00682 | 0.00721 | 0.00762 |

*Note.* Baseline estimates reflect the unperturbed models fitted to digitized data. 5th, 50th (median), and 95th percentiles represent the empirical distribution across 500 stochastic refits under uniform perturbation noise.

**Table S3.**
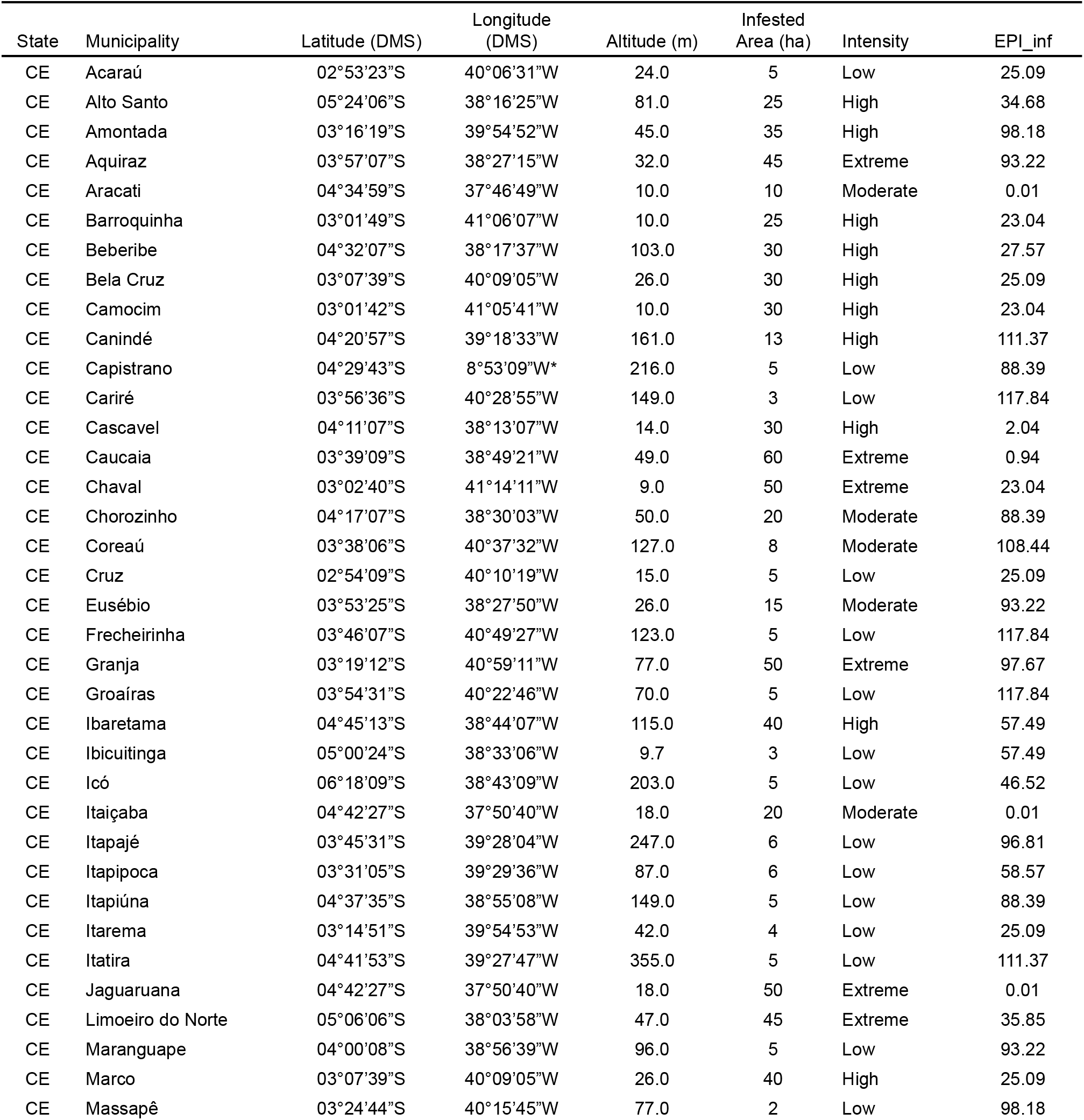
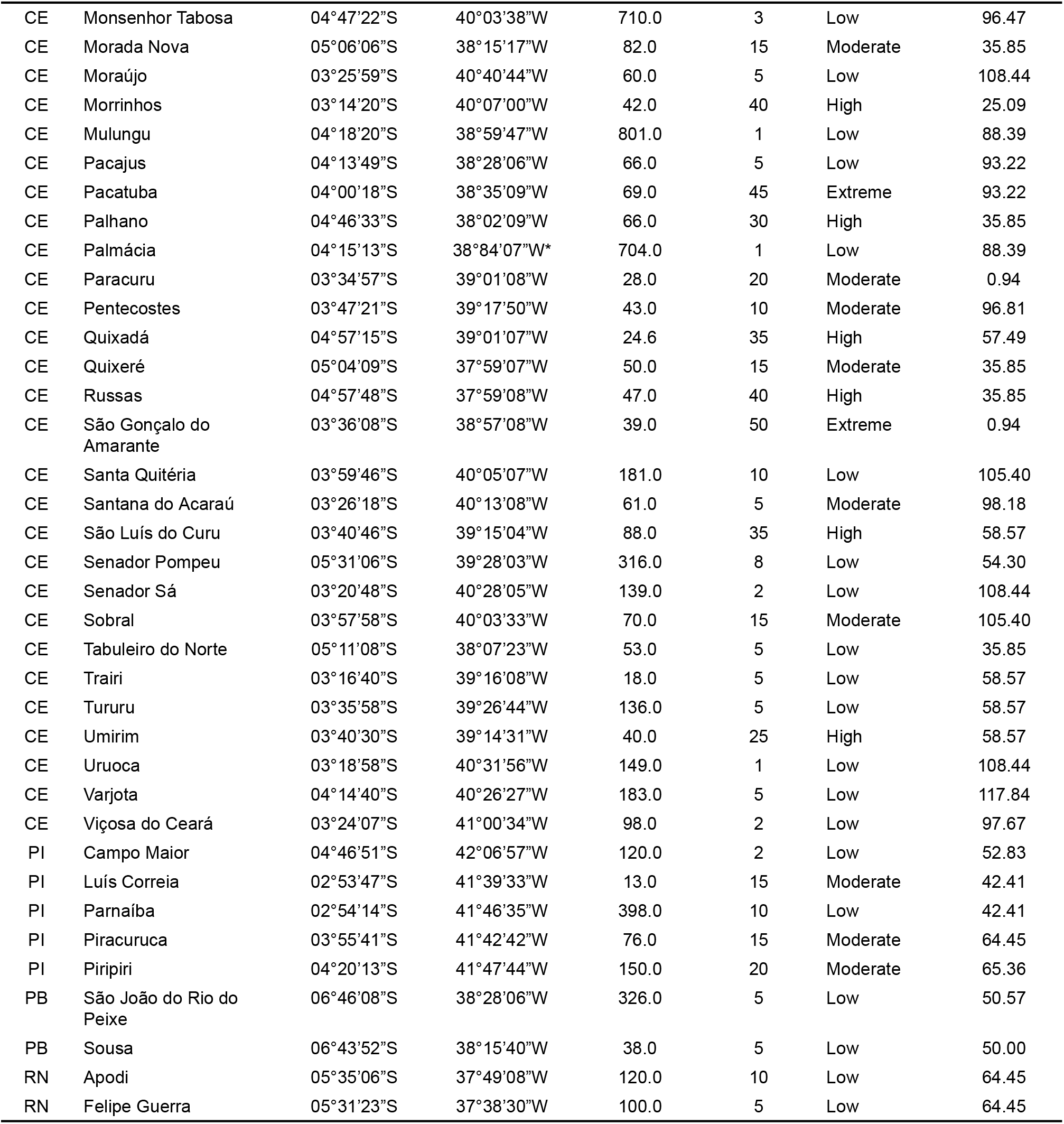
Comprehensive summary of Cryptostegia madagascariensis infestation localities in Northeastern Brazil (reproducing Table 1 from Bonilla et al. 2023) paired with predicted 25-year mean annual weather-driven infection pressure (EPI_inf). *Indicates typographical coordinate errors in the original source publication (Capistrano was published with longitude 8°53’09”W instead of 38°53’09”W, and Palmácia with 38°84’07”W with invalid minute value 84); EPI values were assigned using verified municipal coordinates.

**Figure S1.**
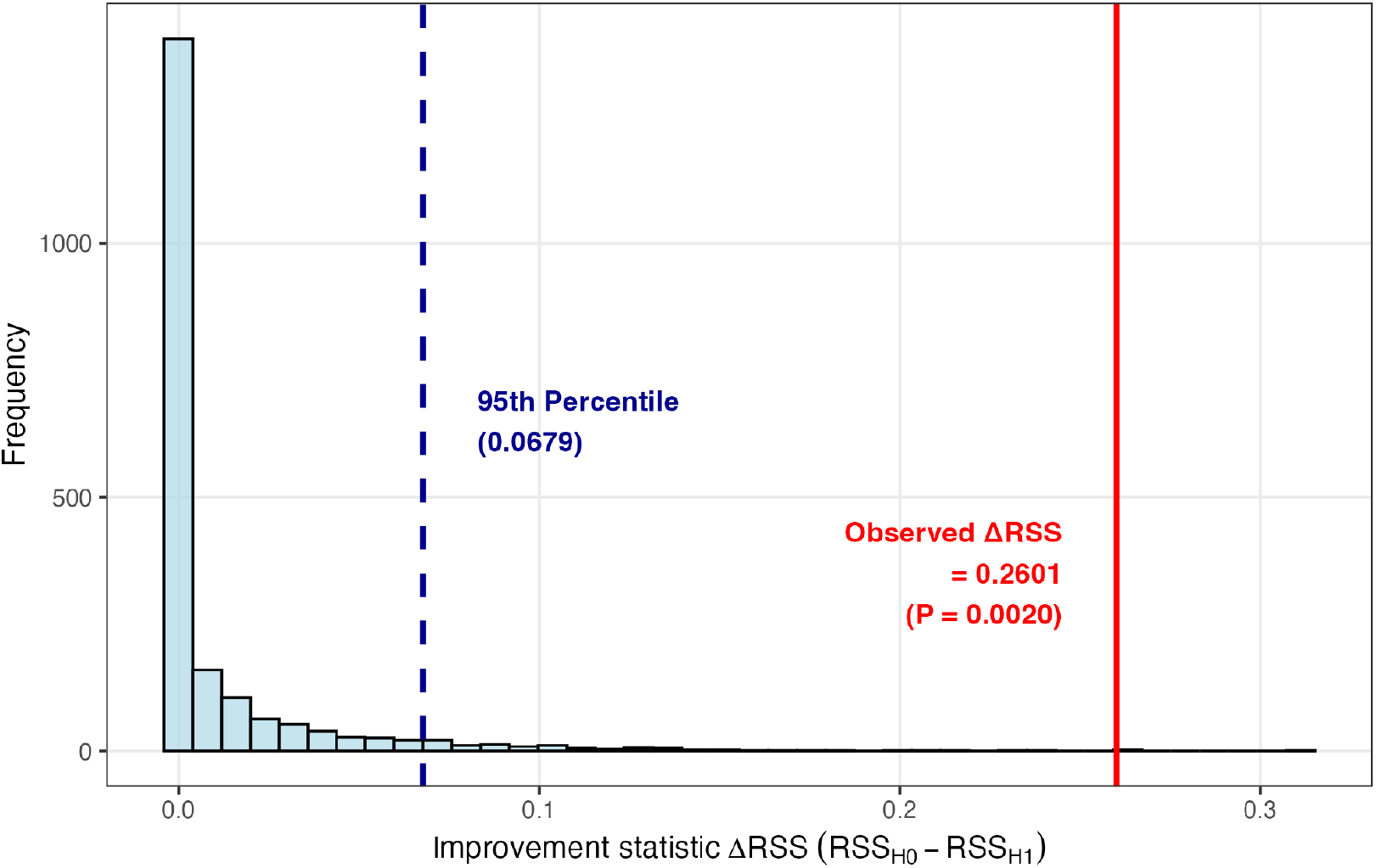
Parametric bootstrap distribution of the improvement statistic (Δ*RSS*) for testing dynamic vs. fixed thermal breadth (γ = 0). The light-blue histogram depicts the empirical distribution of 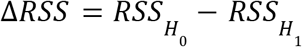 across *B* = 2, 000 parametric bootstrap replicates generated under the null model (*H*_0_: γ = 0; fixed thermal breadth). The vertical dashed navy line indicates the 95th percentile of the null distribution (0.068). The solid red vertical line denotes the observed test statistic (ΔRSS_obs_ = 0.2601, P = 0.002).]

